# Parenteral Vaccination with recombinant EtpA glycoprotein impairs enterotoxigenic *E. coli* colonization

**DOI:** 10.1101/2025.01.03.631264

**Authors:** Tim J Vickers, David Buckley, Nazia Khatoon, Alaullah Sheikh, Bipul Setu, Zachary T. Berndsen, James M. Fleckenstein

## Abstract

Enterotoxigenic *E. coli* (ETEC) cause hundreds of millions of cases of acute diarrheal illness in low-middle income regions, disproportionately in young children. To date there is no licensed, broadly protective vaccine to protect against these common but antigenically heterogeneous pathogens. One of the more highly conserved antigens of ETEC, EtpA, is an extracellular glycoprotein adhesin that preferentially binds to blood group A glycans on intestinal epithelia. EtpA contributes to increased severity of illness in blood group A individuals, elicits robust serologic and fecal antibody responses following infection, and has been associated with protection against subsequent infection. However, its utility as a protective antigen needs further examination. In the present studies we examined whether parenteral vaccination with recombinant EtpA (rEtpA) could afford protection against intestinal colonization in a murine model of ETEC infection. Here, we demonstrate that intramuscular vaccination with rEtpA when adjuvanted with double mutant LT (dmLT) primes IgG predominant mucosal antibody responses to ETEC challenge. Notably, however, both antibody levels and avidity, as well as protection were dependent on vaccination schedule. Likewise, by electron microscopy polyclonal epitope mapping (EMPEM) we observed a greater diversity of epitopes targeted by antibodies after a more protracted vaccination schedule. Next, we explored the utility of IM immunization with alum-adjuvanted rEtpA. This elicited strong serologic and fecal IgG responses. Although accompanied by negligible IgA mucosal responses, EtpA alum-adjuvanted IM vaccination nevertheless protected against ETEC intestinal colonization. Collectively, these data suggest that EtpA could expand the portfolio of antigens targeted in ETEC subunit vaccine development.

## introduction

Enterotoxigenic *Escherichia coli* (ETEC) are a heterogeneous diarrheagenic pathogenic variant (pathovar) of *E. coli* defined by the production of heat-labile and/or heat-stable enterotoxins(1). These pathogens are a leading cause of diarrheal illness in young children in low-middle income countries (LMICs)(2–4) as well as immunologically naïve travelers to areas where clean water and sanitation remain limited(5–12). These pathogens remain a significant cause of death(13) due to acute diarrheal illness among children less than five years of age(2).

Although the death rate from acute illness appears to have declined appreciably over the last several decades due to the implementation of oral rehydration therapy and other measures, infections with these pathogens continue largely unabated(14–16), and have been linked repeatedly to a number of non-diarrheal sequelae including, micronutrient deficiencies(17), malnutrition, and growth impairment(3, 18–21). In addition, malnourished children are at substantially increased risk of death due to diarrhea caused by ETEC and other pathogens(22–24). Underlying the long-term morbidity associated with these pathogens is subclinical intestinal damage characterized by alterations in the absorptive architecture of the small intestine (25), a condition known as environmental enteropathy or enteric dysfunction(26–28).

Most ETEC vaccine development has focused on a group of pathovar-specific plasmid encoded antigens known as colonization factors (CFs). To date, at least 29 distinct antigens have been described (29). The ideal ETEC vaccine, oral or parenteral, would be broadly protective, prevent moderate to severe diarrhea among young children in LMICs during periods when they are most susceptible (6-24 months of age), and be easily integrated for co-administration with available vaccines(30).

The majority of enteric vaccines developed to date, including the most advanced ETEC vaccine to enter clinical trials (31) have relied on oral delivery(32). Notably, however children with enteropathy tend to respond poorly to oral vaccines while their responses to parenterally delivered vaccines remain unimpaired (33, 34). Parenteral subunit vaccination has been shown to induce effective mucosal immune responses (33, 35, 36), and could potentially overcome limitations inherent to oral vaccination in LMICs. Development of fimbrial tip adhesin molecules representing some of the more common CF antigens has provided precedent for pursuit of a parenteral subunit vaccine strategy(37, 38). However, given the heterogeneity of these antigens, it appears likely that other molecules will need to be targeted to achieve broad protection.

One possible candidate antigen, EtpA, is a high molecular weight extracellular ETEC glycoprotein adhesin (39, 40) secreted by a wide variety of enterotoxigenic *E. coli* including some without a recognized colonization factor(41). Molecular epidemiology studies suggest that EtpA-expressing strains are widely geographically distributed(42), and birth cohort studies indicate that this molecule contributes to symptomatic illness in young children(43). EtpA is recognized during natural and experimental(44, 45) human ETEC infections, and prior antibodies to EtpA appear to be associated with decreased risk for subsequent infections(43).

EtpA, like many bacterial adhesins(46), is a lectin. It specifically facilitates ETEC interactions with A blood group glycans on intestinal epithelial cells to promote bacterial adhesion and toxin delivery (47). Intriguingly, children(18) as well as adult human volunteers(47) expressing A blood group antigens appear to be at increased risk of symptomatic ETEC infection. Recent structure-function studies of EtpA, and determination of the molecular structure by cryo-EM, combined with epitope mapping of anti-EtpA monoclonal antibodies have demonstrated that a series of repeat modules comprising the C-terminal region of the molecule direct critical interactions with A blood group glycans to mediate enterocyte adhesion(48).

Elucidation of the nature of these EtpA interactions with the host could be relevant to optimizing its use as a candidate subunit vaccine antigen. To date, recombinant EtpA vaccination via intranasal(49–51), oral, and sublingual(52) routes has been associated with protection against intestinal colonization in a murine model. To explore the utility of EtpA as a parenterally administered antigen subunit, we vaccinated mice intramuscularly with EtpA, examined serologic and mucosal responses to vaccination, and protection against infection with ETEC. Here we demonstrate that EtpA is significantly immunogenic when delivered via IM vaccination, although antibody maturation as well as protection are dependent on timing of immunizations.

## Materials and Methods

### recombinant EtpA production

Recombinant EtpA glycoprotein was produced as previously described(53). Briefly, Top10(pJL017,pJL030) (Table 1) was grown from frozen glycerol stocks maintained at -80°C overnight in 75 ml terrific broth containing carbenicillin (100 µg/ml), chloramphenicol (25 µg/ml), 0.2% glucose, at 37°C, 225 RPM. Overnight growth was then diluted the following morning 1:100 in 2-liter flasks containing 500 ml of fresh media. After growth to OD_600_ of ∼0.6, expression was induced with arabinose (final concentration 0.0002%) for 4.5 hours. Supernatant was recovered by centrifugation at 11,000 x g for 10 minutes, filtered through a 0.2 µm filter, then concentrated by tangential flow (Pellicon 2 Biomax, 100 kDa MWCO) to ∼100 ml. rEtpA was then captured on 2 x 5 ml metal affinity chromatography columns (HisTrap HP, Cytiva Life Sciences), and washed with 5 column volumes of binding buffer (50 mM PO_4_, 300 mM NaCl, pH 7.5). Endotoxin depletion was integrated into subsequent washes on the column with ∼20 column volumes of freshly-prepared 0.1 % (v/v) Triton X-114 (Sigma) in cold PBS(54). The column was then washed with binding buffer (∼10 column volumes) until A_280_ was returned to baseline. Recombinant polyhistidine-tagged EtpA was eluted over a gradient of elution buffer (Elution buffer is 50 mM PO_4_, 300 mM NaCl, 1 M imidazole, pH 7.5). EtpA-containing fractions were identified by SDS-PAGE, pooled and dialyzed vs 10 mM MES, 100 mM NaCl, 1 mM EDTA, pH 6), and concentrated to final concentration of ∼1 mg/ml. Endotoxin levels (EU) were then determined spectrophotometrically (Endosafe-nexgen-PTS, Charles River). Lots used in vaccination were determined to have levels < 50 EU/ml.

**Table 1.**
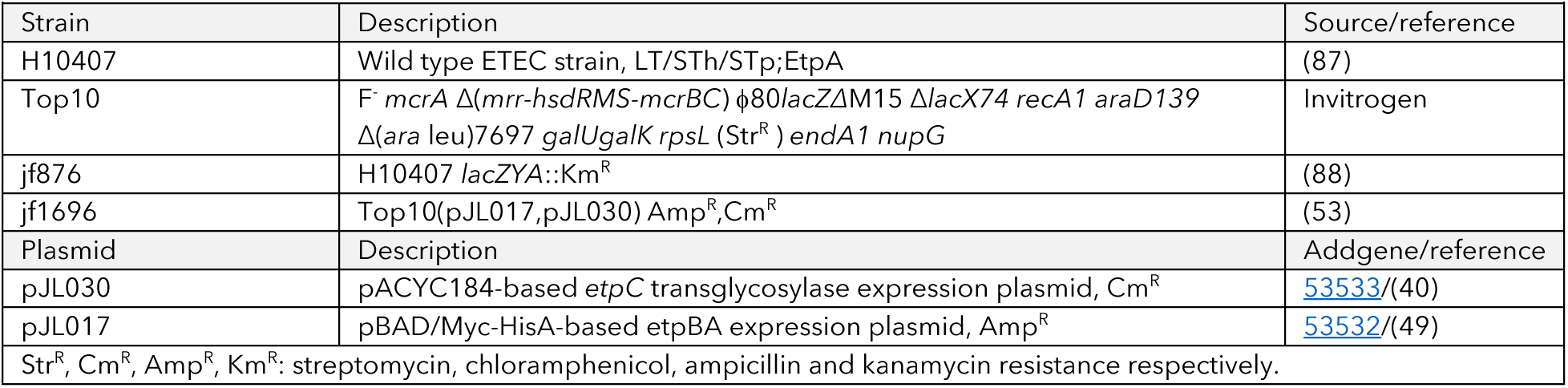
Bacterial strains and plasmids used in these studies.

**Table 2.** Antibodies used in these studies.

| Antibodies used in these studies |  |  |  |  |
| --- | --- | --- | --- | --- |
| Antibody | description | source | Catalog no. | RRID |
| Anti-A blood group | Mouse monoclonal IgM (mAb Z2a) raised vs human A blood group antigen | Santa Cruz Biotechnology | 69951 | AB_1119619 |
| Anti-mouse IgG | Polyclonal horse anti-mouse IgG (H&L) affinity-purified | Cell Signaling | 7076 | AB_330924 |
| Anti-mouse | AlexaFluor 647 conjugated goat anti-mouse IgM heavy chain | Molecular Probes | A21238 | AB_2535807 |
| Anti-O78 | Rabbit anti-O78 polyclonal antisera | Penn State *E. coli* Reference Center | - | - |
| Anti-Rabbit | Goat anti-rabbit IgG (H&L). Alexa Fluor 488 conjugate | Invitrogen | A32731 | AB_2633280 |
| Anti-Rabbit | Goat anti-rabbit IgG (H&L). Alexa Fluor 594 conjugate | Invitrogen | A11012 | AB_2534079 |

### EtpA biotinylation

rEtpA in 10 mM MES (2-(N-Morpholino)ethanesulfonic acid hydrate, Sigma <u>M8250</u>), 100 mM NaCl, pH 6.0 was reacted at room temperature for 30 minutes with a 10 fold molar excess of NHS-LC-LC-biotin (ThermoFisher Sccientific <u>21338</u>) per manufacturer’s directions. The reaction was then quenched with Tris (100 mM, pH 8.0), followed by dialysis in MES buffer to remove free biotin. Biotin incorporation was then confirmed by immunoblotting with streptavidin-HRP.

### ELISA

#### EtpA ELISA

rEtpA, 1 µg/ml in Carbonate buffer, pH 9.6 was used coat wells (100 µl/well) overnight at 4° C, then washed 3 x with 200 µl of PBS-0.05% Tween-20 (PBS-T). Plates were blocked (1 h, 37° C) by incubation with 200 µl/well of 1% BSA in PBS-T. Primary sera were diluted in PBS-T-BSA as indicated, and 100 µl was added to respective wells for 1 hour at 37° C. Wells were washed 5x with PBS-T, then horse anti-mouse IgG -HRP conjugate 2° antibody (Cell Signaling <u>7076</u>) added in PBS-T-BSA, 100 µl/well, 37°C, x 1 hour. Wells again washed 5x with PBS-T and detected with freshly prepared TMB peroxidase substrate (3,5,3’,5’-tetramethylbenzidine, seracare <u>5120-0053</u>), then read kinetically(55) at 650 nm (Eon, Biotek).

#### IgG avidity assay

ELISA plates were coated with rEtpA (1.0 µg/ml) in carbonate buffer (pH 9.6) at 4°C overnight. Plates were then washed 6x with PBS-0.05 % Tween-20, and blocked with 1 % BSA in PBS-0.05 % Tween-20 for 1 hour at 37°C. Sera diluted 1:100,000 in blocking buffer were incubated for one hour at 37°C, then washed with PBS. Half of the plate was incubated for an additional 5 minutes in PBS while the other half was treated for 5 min with 8M urea in PBS, After washing, affinity purified horse anti-mouse IgG conjugated to HRP (Cell Signaling <u>7076</u>) diluted 1:5000 in PBS-T-BSA was added, followed by incubation for 1 hour at 37°C. Plates were then washed with PBS and developed with freshly prepared TMB substrate. Antibody avidity was calculated as the ratio of kinetic ELISA responses in urea-treated wells relative to corresponding untreated wells (56).

#### Blood group A-EtpA binding ELISA

To examine EtpA binding to human A blood group (bgA), BSA-conjugated blood group A (MO Bi Tec Dextra <u>NGP6305</u>) was dissolved to a final concentration of 0.5 mg/ml in PBS containing 0.02 %(w/v) azide, and stored at 4° C immediately prior to use. bgA-BSA conjugate working solution was then prepared in carbonate buffer, pH 9.6 (final concentration of 1 µg/ml). ELISA strips (Corning 2580) were incubated overnight (100 µl/well) at 4° C, then washed 3 x with 200 µl of PBS containing 0.02 % Tween-20 (PBS-T), and blocked with 100 µl of 1 % BSA in PBS-T at 37° C for 1 hour. 100 µl of biotinylated EtpA-biotin (10 µg/ml in PBS-T-1 % BSA) was added per well, incubated for 2 hours at room temperature, then washed 5 x with of PBS-T. Wells were then incubated for 1 hour with avidin-HRP conjugate (BioRad 1706528, diluted 1:10,000 in 1 % BSA in PBS-T) at room temperature, then washed again 4x with PBS-T. Finally, wells were developed with freshly-prepared room-temperature HRP substrate (TMB-(3,5,3’,5’-tetramethylbenzidine)-2 component reagent (<u>seracare 5120-0053</u>), and read kinetically at 650 nm.

### Intramuscular vaccination

Mice were vaccinated intramuscularly (IM) with 10 µg of rEtpA with 1 µg of double (R192G/L211A)(57) mutant LT (dmLT) in a final volume of 50 µl. Control mice were vaccinated with 1 µg of dmLT alone, or an equal volume of PBS. Mice were vaccinated in two immunization schedules on days 0, 14, 28, or days 0, 21, 42. Alternatively, mice were vaccinated on days 0, 21, 42 with aluminum phosphate adjuvant alone (Adju-Phos <u>vac-phos-250</u>, InvivoGen, San Diego, CA, USA), or 10 µg of rEtpA adjuvanted with Adju-Phos in a volume-volume ratio of 1:1.

### Murine Small intestinal colonization

Colonization studies were carried out in streptomycin-treated adult CD-1 mice as previously described(52). Strain jf876 was grown overnight in kanamycin (25 µg/ml) in 2 ml of LB at 37°C, 225 rpm, diluted in fresh media the morning of challenge and grown to OD600 of ∼0.3. Mice were challenged by gavage with ∼10^5^ colony forming units. On the day after challenge small intestinal segments were isolated, lysed in saponin (5 %) for 10 minutes and serial dilutions plated onto Luria agar plates containing kanamycin. All experiments were conducted with the approval of the Animal Care and Use Committee at Washington University School of Medicine.

### ETEC adhesion to blood group A intestinal epithelia

HT-29 cells (<u>ATCC HTB-38</u>), which express blood group A glycans (47), were propagated as previously described in McCoy’s-5A medium (Gibco, Life Technologies) supplemented with 10 % bovine serum albumin. Cells were grown to confluence in 96-well plates at 37° C, 5 % CO_2_. Adhesion assays were performed as previously described(47) using mid-log phase bacterial cultures. After 30 minutes monolayers were washed 3 x with pre-warmed media, then treated with 0.1 % Triton-X-100 in PBS for five minutes. Dilutions of the resulting lysates were plated onto Luria agar and bacterial adherence expressed as the percentage of the original inoculum recovered. Alternatively, adherent bacteria were identified by confocal microscopy.

### Confocal laser scanning microscopy

HT-29 cells seeded onto pre-treated poly-L-lysine glass coverslips in 24 well plates were incubated and grown as above at 5% CO_2_, 37°C to confluence. ETEC H10407 was added at a multiplicity of infection of ∼1:100 and incubated for ∼30 minutes prior to fixation. Plasma membranes were stained with CellMask deep red (<u>Thermo Fisher Scientific, C10046</u>) (1:2,000) and nuclei with DAPI (1:6000). BgA was detected with mouse monoclonal antibody Z2A (<u>Santa Cruz sc-69951</u>) against human A blood group antigen, followed by AlexaFluor 647-conjugated goat anti-mouse IgM heavy chain (<u>Molecular Probes, A21238</u>). Confocal images were acquired using a Nikon Eclipse Ti2 inverted microscope. ETEC H10407 (serotype O78) were imaged using polyclonal antisera (Rabbit) supplied by the Penn State E. coli Reference Center, followed by cross-absorbed goat anti-rabbit IgG (H\&\L) conjugated to either AlexaFluor 488 or 594 fluorophores (Invitrogen).

### Electron Microscopy

#### Negative Stain Electron Microscopy Polyclonal Epitope Mapping (nsEMPEM) sample preparation

Samples of ∼ 1mg/mL IgG purified from sera of mice vaccinated with rEtpA/dmLT at 2 or 3 week intervals were incubated with 0.5 mL of immobilized papain resin (Thermo Fisher Scientific) for 6 h at 37°C to liberate Fab from Fc regions the antibodies. The digestion mixtures of 2-week and 3-week polyclonal samples were separated from the papain resin by centrifugation at 4,698 x g for 10 min. Size-exclusion chromatography (SEC) was performed on each isolated digestion mixture using a Superdex 200 Increase 10/300 GL column (Cytiva), and fractions from the Fab/Fc-containing peak were pooled. Pooled 2-week and 3-week samples were buffer-exchanged in parallel into the rEtpA suspension buffer (10 mM MES, 100 mM NaCl, pH 6.0) and subsequently concentrated to ∼0.1 mg/mL using 10 kDa-cutoff Amicon Ultra Centrifugal filter units (Sigma-Aldrich). Fab-rEtpA complexes were formed by incubating a 6x molar excess of each purified Fab/Fc 2-week and 3-week sample with 15 µg rEtpA to a final volume of 300 µL and stored at 4°C for 36 h. SEC was repeated on the incubated samples to separate rEtpA-Fab complexes from free rEtpA/Fab/Fc.

##### Negative stain Electron microscopy (nsEM) data acquisition

Freshly purified 2-week and 3-week rEtpA-Fab complexes were concentrated to ∼0.05 mg/mL. For each sample, 3 µL was pipetted onto glow-discharged carbon-coated 300-mesh Cu grids, and immediately blotted with filter paper. After repeated pipetting and blotting (x 2), grids were negatively stained by incubating 3 µL of 2% (w/v) uranyl acetate for 30 seconds. Data were collected on the 120keV JEOL JEM-1400Plus electron microscope at the Washington University Center for Cellular Imaging (WUCCI). A magnification of x30,000 was used throughout data collection, with a nominal pixel size of 3.54 Å. Micrographs were obtained manually, using an AMT XR111 high-speed 4k x 2k pixel phosphor-scintillated 12-bit CCD camera. CryoSPARC v4.6.2(58) was used for all data processing.

##### nsEMPEM data processing

Elliptical blob-picking and subsequent pick filtering was performed on 323 micrographs for the 2-week rEtpA-Fab dataset, resulting in a final set of 2,574 rEtpA-Fab particles (∼9 particles/micrograph) after several rounds of 2-dimensional (2D) classification followed by subset selection. Ten final, representative 2D classes (of 1,575 particles) were obtained for the 2-week dataset after 2D class rebalancing, subset selection, and additional 2D classification.. Manual picking of particles from 934 micrographs for the 3-week rEtpA-Fab dataset was performed, and 3,439 rEtpA-Fab particles (∼4 particles/micrograph) were obtained after several rounds of 2D classification followed by subset selection. 2D classes of manually-curated particles from the 3-week rEtpA-Fab dataset were used to perform a secondary template-based particle picking, which was combined with the original manually-picked particles through the ‘Remove Duplicates’ job following several more rounds of 2D classification and subset selection (resulting in a final total particle count of 4,956). Twenty final, representative 2D classes were split into two sets of ten 2D classes that highlight multiple Fabs bound to EtpA (1,338 particles), and single Fabs bound to EtpA (1,527 particles) through 2D class rebalancing and subsequent rounds of 2D classification.

## results

### Parenteral vaccination primes mucosal responses to EtpA

A number of prior studies have demonstrated that parenteral vaccination, when adjuvanted with ADP-ribosylating enterotoxins, including the modified double (R192G/L211A) mutant version *E. coli* heat-labile toxin (dmLT)(59), can direct antigen-specific immune responses in the intestine(60–62). To explore the utility of parenteral vaccination with EtpA, we first questioned whether IM vaccination of rEtpA adjuvanted with dmLT would engender EtpA-specific mucosal antibody responses. Intramuscular vaccination of mice resulted in fecal IgA as well as IgG responses to EtpA, with fecal IgG being predominant (figure 1A). Notably, both IgA and IgG mucosal responses increased significantly following ETEC infection of parenterally immunized mice relative to PBS controls where no response to infection alone was observed. These data suggested that parenteral vaccination with rEtpA, at least when adjuvanted with dmLT, can elicit mucosal antibody responses and that this strategy may prime mucosal responses to this antigen elicited by ETEC infection.

**Figure 1.**
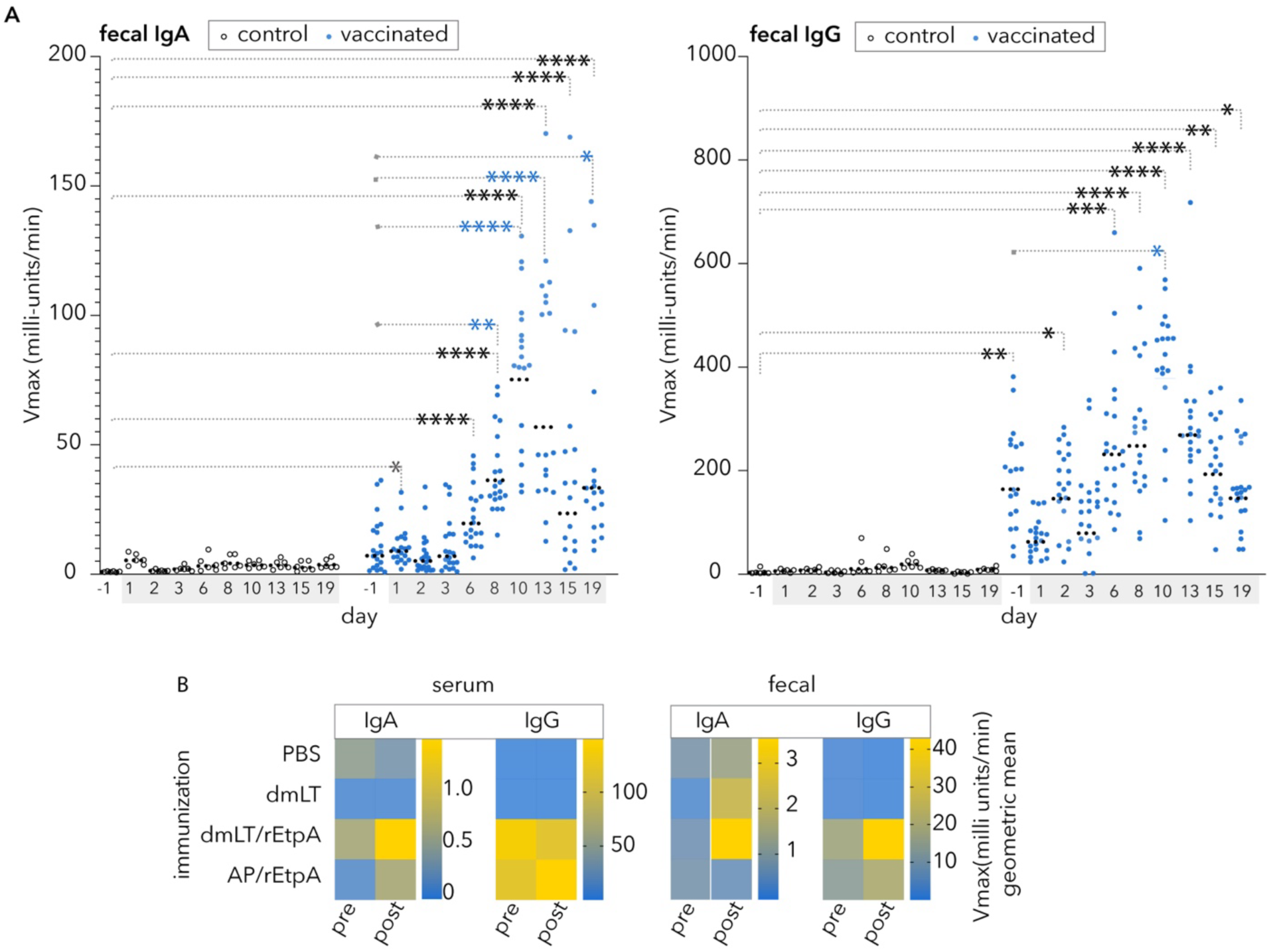
Parenteral vaccination with recombinant EtpA (rEtpA) adjuvanted with dmLT elicits mucosal antibody responses and enhances antigen responses following infection. **A.** Graphs depict anti-EtpA fecal antibody responses by kinetic ELISA for sham-vaccinated (PBS, open circles) controls (n=6), and mice vaccinated with rEtpA adjuvanted with dmLT (blue symbols, n=20), before (day -1) and after (shaded days) infection with ETEC H10407 (1.5 x 10^7^ cfu). Fecal resuspensions were tested at 1:4 for both isotypes. Comparisons by Kruskal-Wallis testing: *<0.05, **<0.01, ***<0.001, ****<0.0001. Significant comparisons to PBS d-1 controls and vaccinated d-1 are shown in black and blue, respectively. Dashed horizonal lines represent geometric means. **B.** Heatmaps summarize kinetic ELISA data (geometric mean values) following vaccination of groups of mice (n=5) with PBS (sham), dmLT adjuvant alone, rEtpA adjuvanted with dmLT (dmLT/rEtpA), or rEtpA adjuvanted with AdjuPhos (AP). Sera were tested at 1:500 dilution (IgA) and 1:100,000 (IgG). Fecal samples were assessed at a 1:10 dilution. Post challenge samples were obtained on day 8 following infection with ETEC H10407 (2 x 10^5^ cfu).

### Comparison of dmLT to alum

We next compared both serum and fecal responses to EtpA following parenteral vaccination with dmLT as the adjuvant relative to vaccination with rEtpA adjuvanted with alum. Vaccination with either adjuvant preparation resulted in modest serum IgA responses but robust IgG responses (figure 1B). We observed modest increases in fecal IgA following H10407 challenge in mice immunized with dmLT/rEtpA, but not in mice in which alum was used at the adjuvant. Anti-EtpA fecal IgG levels were increased in both groups, although somewhat higher in mice immunized with the dmLT-EtpA combination.

### Timing of rEtpA parenteral vaccination adjuvanted with dmLT dictates outcome

Next, we examined the protection afforded against ETEC small intestinal colonization by parenteral immunization in a murine model of infection(49, 63). Mice were vaccinated IM with 10 µg of rEtpA adjuvanted with 1 µg of dmLT on days 0, 14, 28 and then challenged on day 44 with ∼2 x 10^5^ colony forming units of jf876 to assess protection against ETEC colonization of the small intestine. We again observed IgG predominant responses in both feces and sera (figure 2A). However, this was not associated with significant protection against intestinal colonization relative to adjuvant-only vaccinated controls (figure 2B).

**Figure 2.**
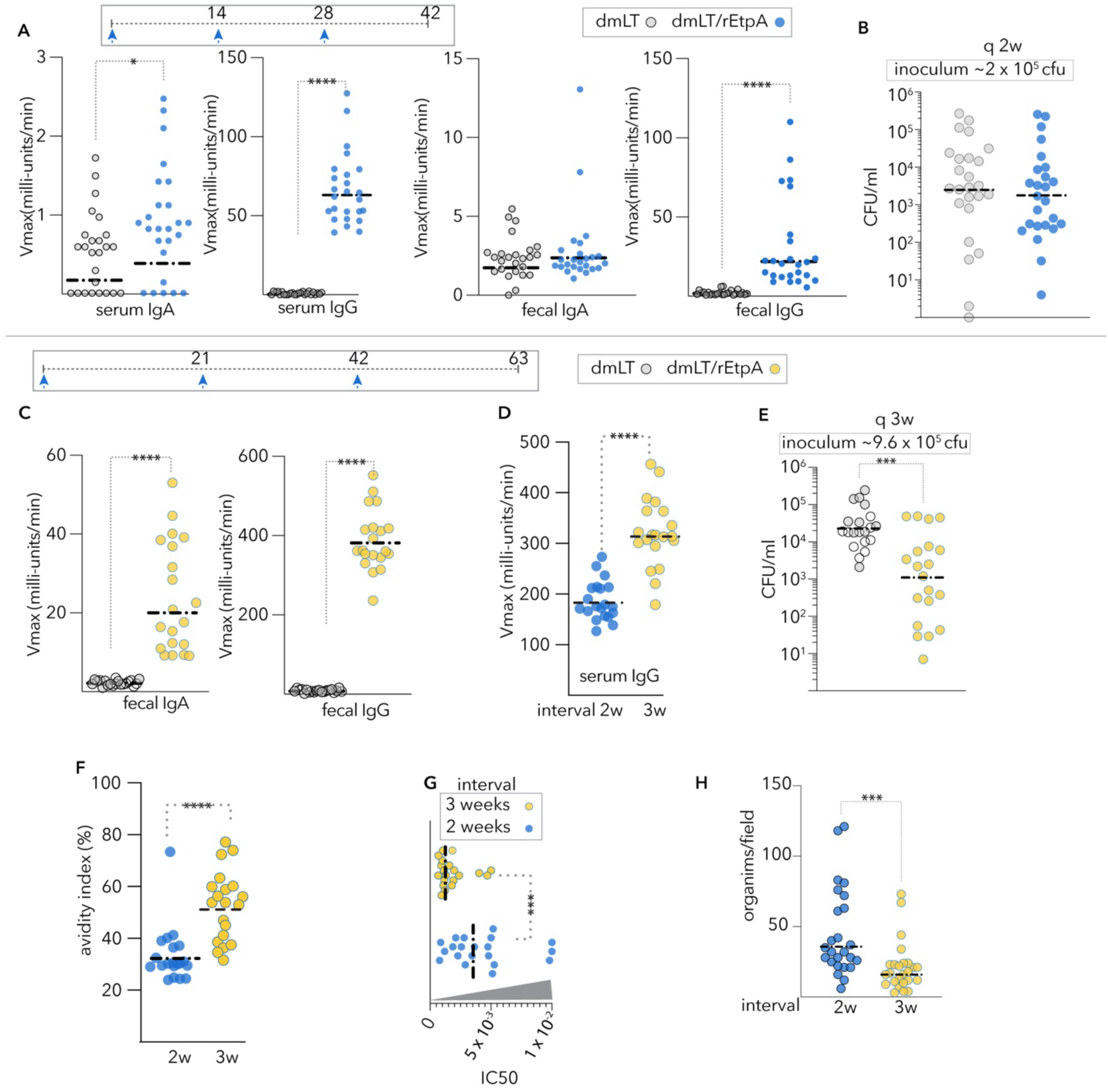
Vaccination timing impacts immunogenicity and protection. **A.** Serum and Fecal IgA and IgG values following IM immunization on days 0,14,28 with rEtpA adjuvanted with dmLT. **B.** Intestinal colonization at day 42 in unvaccinated controls vs mice vaccinated with rEtpA adjuvanted with dmLT at 2-week intervals. **C**. anti-EtpA fecal IgA, and IgA following IM vaccination at 3-week intervals with dmLT/rEtpA vs unvaccinated controls anti-EtpA fecal IgG following IM vaccination at 3-week intervals with dmLT/rEtpA vs unvaccinated controls **D.** Comparison of anti-EtpA serum IgG values (kinetic ELISA) of mice vaccinated with rEtpA/dmLT at 2- or 3-week intervals. (sera diluted 1:100,000). **E**. Intestinal colonization in unvaccinated controls vs mice vaccinated with rEtpA adjuvanted with dmLT at 3-week intervals. **F**. Avidity of IgG antibody following vaccination with dmLT/rEtpA at 2- or 3-week intervals. **G**. Vaccination at 3-week intervals enhances neutralization activity of IgG antibody in rEtpA-blood group A binding assays. **H.** IgG from mice vaccinated at 3-week intervals more effectively impairs ETEC adhesion to target HT29 cells. Dashed horizontal bars throughout represent geometric mean values. Statistical comparisons between groups by Mann-Whitney (2-tailed) analysis. ****<0.0001, ***<0.001,*<0.05.

Because the pace of antigen delivery can influence B cell maturation and antibody affinity as well as the diversity of epitopes recognized (64, 65) we vaccinated a second group of animals with the same adjuvant and dose of rEtpA, but at 3 week intervals (days 0, 21, 42, followed by challenge on day 63). Again, we observed a predominant fecal anti-EtpA IgG response to vaccination (figure 2C). Interestingly, mice vaccinated with rEtpA adjuvanted with dmLT on this more protracted schedule generally exhibited stronger serum IgG responses than those vaccinated on a every 2 week schedule (figure 2D), and were protected against intestinal colonization with ETEC relative to adjuvant only controls (figure 2E). Finally, the avidity of serum antibodies was appreciably higher (figure 2F) following vaccination at 3 week intervals, and these antibodies were also significantly more effective in preventing EtpA interaction with target blood group A molecules (figure 2G), and EtpA-expressing ETEC adhesion to target blood group A + HT29 cells (figure 2H).

Our recent studies have shown that the C-terminal repeat region of EtpA is essential for its activity as a blood group A lectin and that monoclonal antibodies which recognize the repeat regions of EtpA interrupt binding to target glycans (Figure 3a)(48). Therefore, to further characterize humoral responses to rEtpA vaccination we performed negative stain electron microscopy-based polyclonal epitope mapping (EMPEM)(66) of Fab fragments generated from IgG isolated from mice vaccinated with rEtpA adjuvanted with dmLT (figure 3b, supplementary figure 1). We identified a greater diversity of epitopes targeted by Fabs isolated after the every 3 week schedule of as well as evidence of rEtpA antigens with multiple bound Fabs, including Fabs targeting neutralizing epitopes on the C-terminal repeat domain (CTR), which allow for binding avidity of intact IgG(48). Collectively, these results demonstrate that immunization timing can profoundly affect both the quantity as well as quality of antibodies elicited against rEtpA, findings which coincide with the ability to neutralize EtpA lectin activity *in vitro* and ultimately protection against colonization *in vivo*.

**Figure 3.**
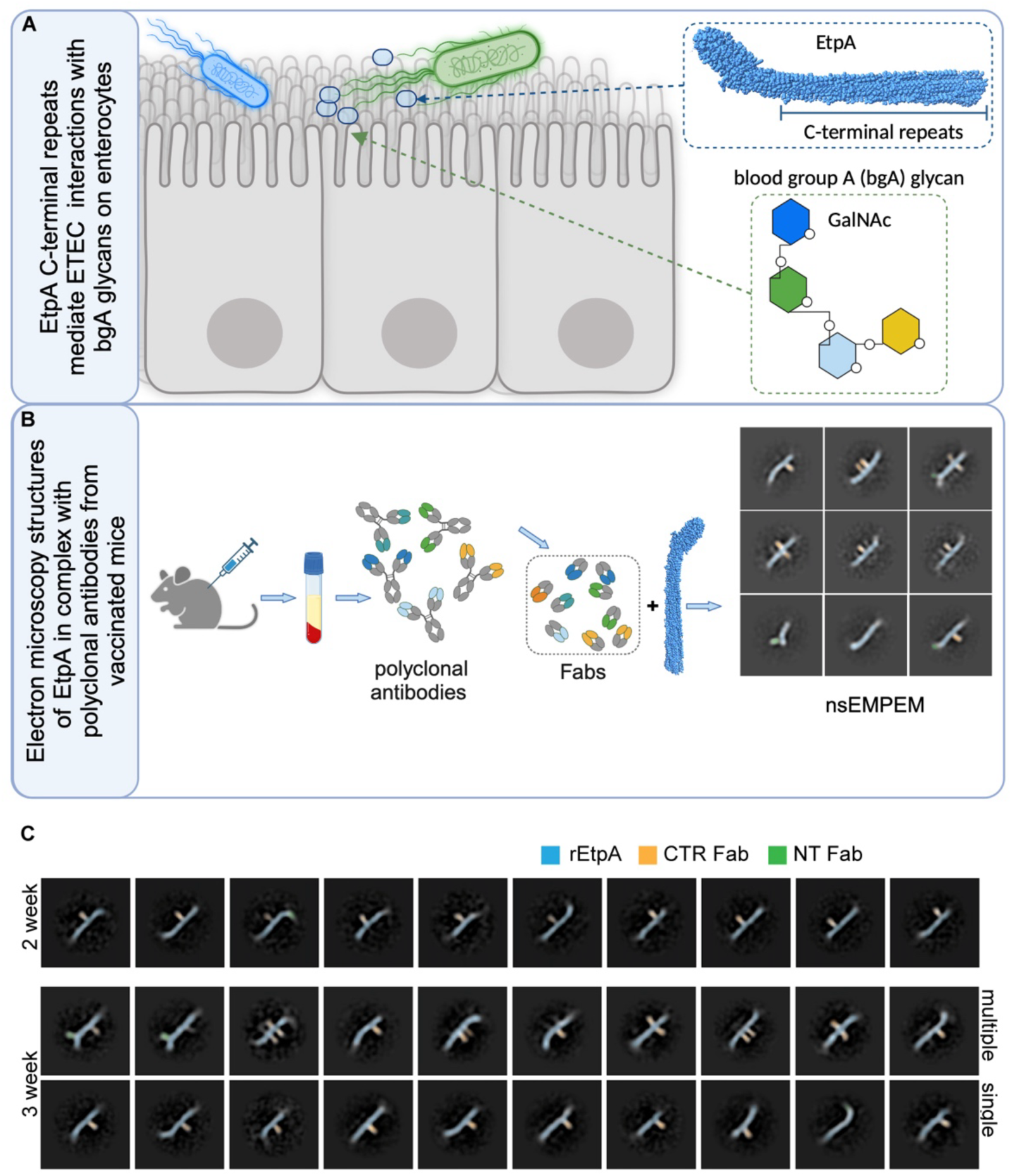
Vaccination administration schedule impacts antibody diversity. **A.** Schematic of EtpA lectin activity wherein C-terminal repeat region of EtpA (CTR) directs binding to blood group A glycans expressed on the surface of enterocytes. **B.** negative stain electron microscopy polyclonal antibody epitope mapping (nsEMPEM) protocol. **C.** nsEMPEM of IgG Fabs generated from sera of mice vaccinated every 2 weeks or every three weeks with rEtpA adjuvanted with dmLT. Selected 2-dimensional class averages of rEtpA in complex with Fabs generated from sera of mice vaccinated every 2 weeks (top row). Middle panel shows highlights multiple-Fabs/rEtpA molecule, and bottom row shows single-Fabs/rEtpA from mice vaccinated every 3 weeks with rEtpA/dmLT. Pseudo-coloring reflects rEtpA (blue), CTR-binding Fabs (orange), and NTD-binding Fabs (green).

### Parenteral vaccination with alum-adjuvanted rEtpA impairs ETEC colonization

Although mutant versions of heat-labile toxin have now been used in multiple clinical trials, including recent subunit ETEC vaccine studies(67), many vaccines licensed over the past several decades have been adjuvanted with alum(68). Therefore, to extend our observations we examined immune responses and protection afforded by vaccination with rEtpA vaccinated with aluminum phosphate following the more protracted vaccination schedule on days 0,21,42 with challenge on day 63. Alum-adjuvanted IM vaccination with rEtpA yielded a predominant IgG response in both serum (figure 4A) and stool (figure 4B). Despite little discernable IgA response, vaccinated mice exhibited reduced levels of intestinal colonization with ETEC following challenge (figure 4C) further suggesting that rEtpA could afford protection as part of a parenterally delivered subunit vaccine.

**Figure 4.**
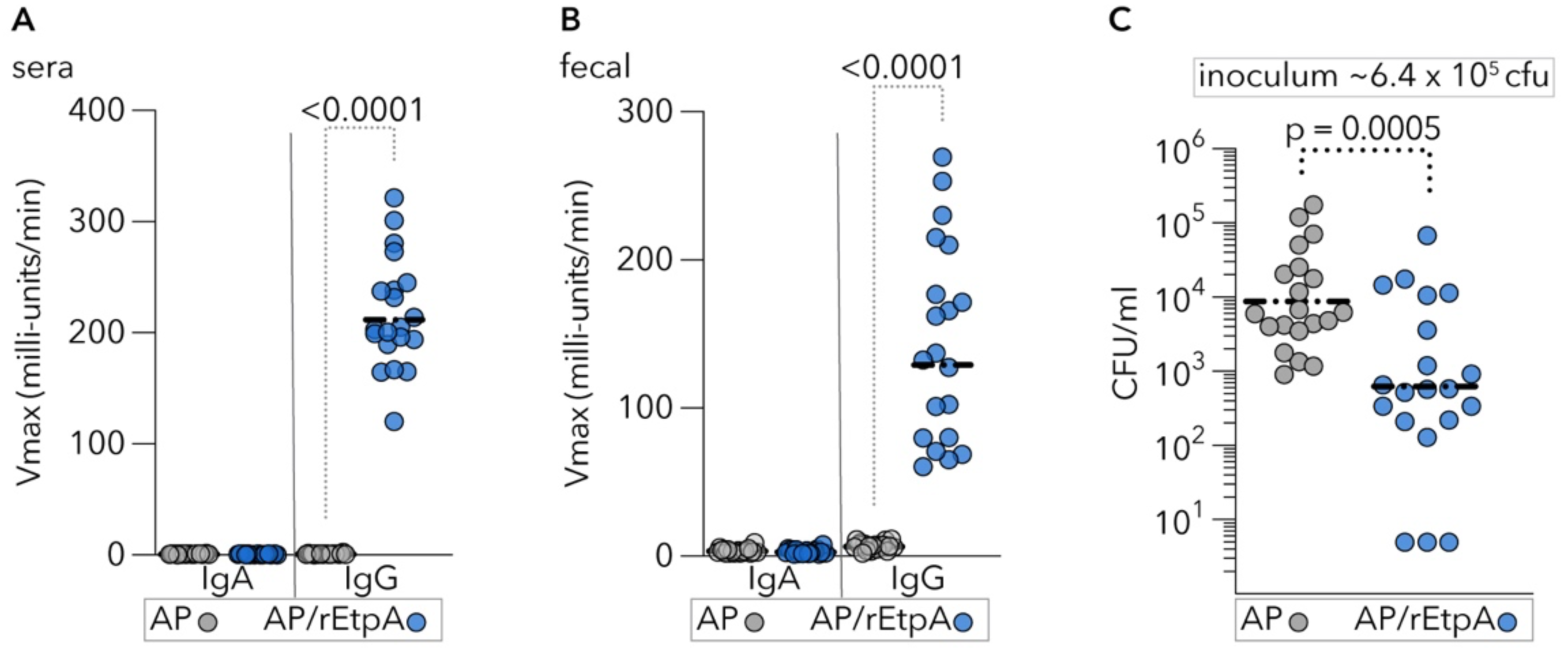
Intramuscular vaccination with alum-adjuvanted EtpA protects against intestinal colonization with ETEC. CD-1 mice (n=20/group) were vaccinated (days 0, 21, 42) with Adju-Phos (AP) alone or EtpA adjuvanted with AP. **A**. Serum IgA and IgG levels following vaccination with AP alone or rEtpA adjuvanted with AP. **B**. anti-EtpA fecal antibody levels following vaccination. **C. Intestinal colonization of** mice following challenge on day 63. Comparisons by Mann-Whitney (2-tailed). Dashed horizontal lines represent geometric mean values.

## Discussion

Since their discovery as a cause of acute cholera-like watery diarrhea more than 50 years ago(69, 70), enterotoxigenic *E. coli* have presented formidable challenges to vaccine development. In contrast to *Vibrio cholerae,* which have largely been limited to only a few serotypes, the plasmid-encoded virulence genes of ETEC are present in a genetically diverse group of *E. coli* comprised of a multitude of O- and H-serogroups(71).

The main targets for vaccine development over the past five decades have been plasmid-encoded antigens known as colonization factors (CFs)(72). Early enthusiasm for CF antigens was in part stimulated by studies demonstrating that a strain (H10407-P) cured of a plasmid carrying genes for the CFA/I colonization factor was effectively avirulent relative to the ETEC wild type H10407 strain(73). Subsequent studies demonstrated that passive immunization with hyperimmune bovine milk immunoglobulin raised against CFA/I afforded significant protection against challenge with ETEC H10407(74) engendering additional support for CF-based ETEC vaccine development. More recently, sophisticated structural studies have facilitated the development of recombinant tip adhesin molecules which likewise offer protection in passive(75) and active(76) immunization studies.

Despite these advances, development of a broadly protective subunit vaccine for ETEC based exclusively on CFs will present significant challenges given the extraordinary diversity of these antigens described to date. Notably, the large mosaic plasmid encoding CFA/I previously cured from H10407 was later shown to encode the etpBAC two-partner secretion locus responsible for production of the EtpA glycoprotein(39), as well as the EatA mucinase autotransporter(77).

It is likely that subunit vaccine development incorporating EtpA will require further exploration and optimization of adjuvants as has been done with fimbrial subunits(78). Nevertheless, the present studies suggest that parenteral immunization with EtpA is feasible, and that this antigen could expand coverage afforded by current CF-centered approaches. Recent studies have demonstrated that non-mucosal vaccination adjuvanted with dmLT can elicit antigen-specific migration of CD4+ antigen-specific T cells(60), and B cells into intestinal mucosa resulting in fecal IgA(79). In the current study, IM vaccination with EtpA adjuvanted by dmLT elicited modest fecal IgA and IgG responses, with demonstrable increases in antigen-specific antibody responses following infection. IgA produced at mucosal sites is thought to exclude enteric pathogens in part by clumping the growing bacteria to accelerate their clearance from the intestine(80).

The majority of parenteral vaccines, including several in the Expanded Program on Immunization, have been adjuvanted with alum(81). Despite a traditional focus on mucosal protection mediated by IgA, IgG has been shown to play a complementary if not dominant role in protection against some intestinal pathogens(82) in part by targeting surface virulence factors triggering elimination by transmigrating neutrophils(83). The studies reported here suggest that the predominant fecal IgG responses generated by immunization with recombinant EtpA adjuvanted with either dmLT or alum may sufficiently target this surface-expressed adhesin to provide protection.

The ability to define precise mechanistic correlates of protection remains a challenge for many vaccines targeting mucosal pathogens(84–86). However, detailed understanding of antigenic structure-function relationships can aid in characterization of neutralizing antibody responses that lead to protection. The recent determination of the EtpA structure and the elucidation of the essential role played by its C-terminal repeat domain in directing interactions with target glycans on mucosal epithelia(48) can facilitate development of assays that help to define its role as a protective subunit antigen and aid the identification of correlates of protection.

## Supplementary information

**Supplementary figure 1.**
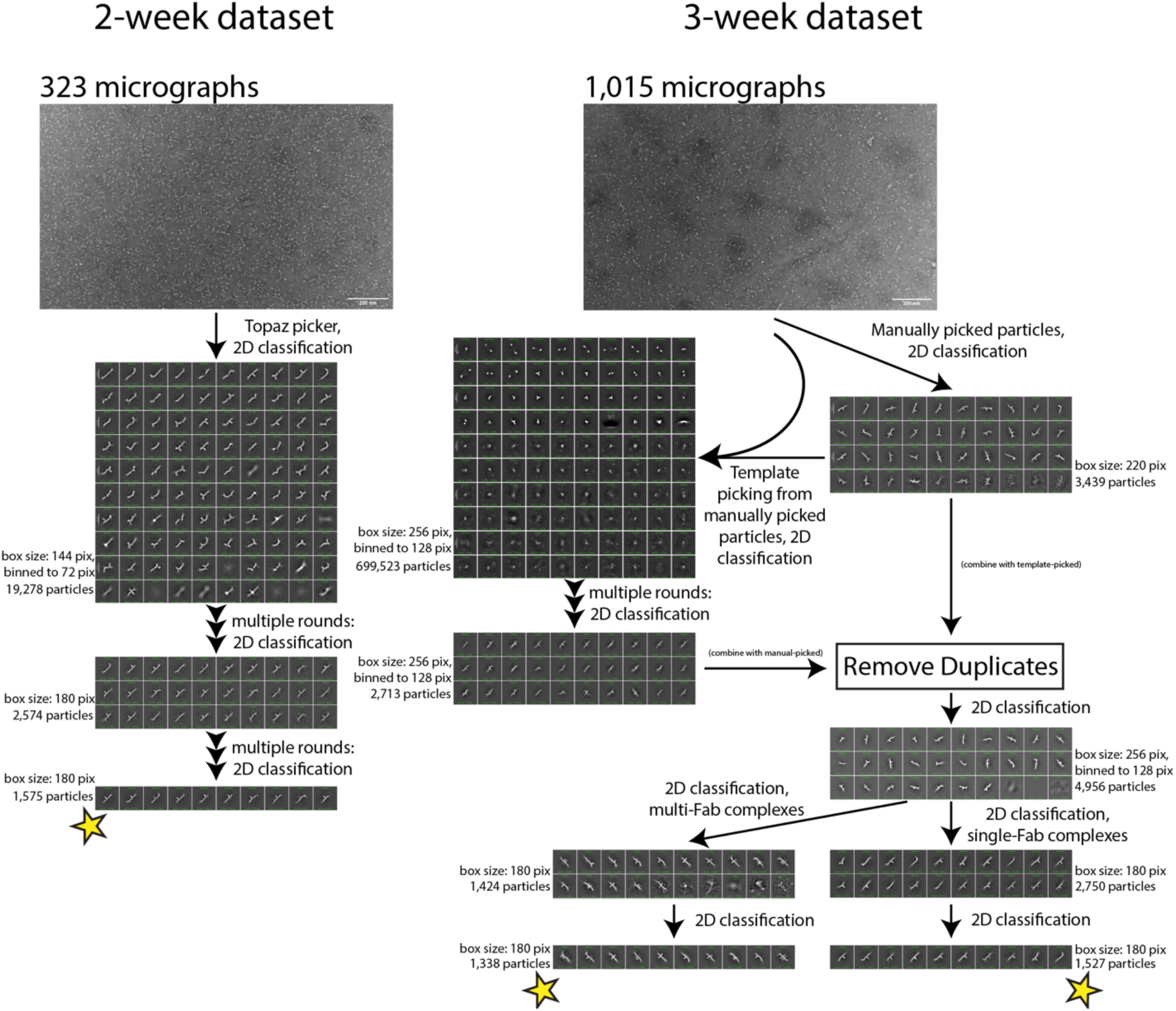
nsEMPEM particle identification strategy.

